# A brain-shuttled antibody targeting alpha synuclein aggregates for the treatment of synucleinopathies

**DOI:** 10.1101/2024.10.16.618734

**Authors:** Sungwon An, John J. McInnis, Dongin Kim, Yihang Li, Ozge Yilmaz, Jinhyung Ahn, Brian C Mackness, Jinyoung Park, Julia Maeve Bonner, Hyesu Yun, Simon Dujardin, Donghwan Kim, Seung-Hwan Kwon, Yi Tang, Laurent Pradier, Sumin Hyeon, Daehae Song, Byungje Sung, Rajaraman Krishnan, Brian Spencer, Robert Rissman, Jagdeep K. Sandhu, Arsalan S. Haqqani, Hyeran Lee, Jinwon Jung, Weon-Kyoo You, Alexandra T. Star, Christie E. Delaney, Danica B. Stanimirovic, Sergio Pablo Sardi, Sang Hoon Lee, Can Kayatekin

**Affiliations:** ABL Bio, Inc.; Sanofi; University of Southern California; University of California San Diego; National Research Council Canada

**Author notes:** Current address: University of Southern California, Alzheimer’s Therapeutic Research Institute (Robert Rissman). Deceased (Danica B. Stanimirovic).

## Abstract

Parkinson’s disease and multiple system atrophy are members of a class of devastating neurodegenerative diseases called synucleinopathies, which are characterized by the presence of alpha-synuclein (α-Syn) rich aggregates in the brains of patients. Passive immunotherapy targeting these aggregates is an attractive disease-modifying strategy. Such an approach must not only demonstrate target selectivity towards α-Syn aggregates, but also achieve appropriate brain exposure to have the desired therapeutic effect. Here we present preclinical data for a next-generation antibody for the treatment of synucleinopathies. SAR446159 (ABL301) is a bispecific antibody composed of an α-Syn-binding immunoglobulin (IgG) and an engineered insulin-like growth factor receptor 1 (IGF1R) binding single-chain variable fragment (scFv), acting as a shuttle to transport an antibody across the blood-brain barrier (BBB). SAR446159 binds tightly and preferentially to α-Syn aggregates and prevents their seeding capacity *in vitro* and *in vivo*. Incubation with SAR446159 reduced α-Syn preformed fibrils (PFFs) uptake in neurons and facilitated uptake and clearance by microglia. In wild type mice injected in the striatum with α-Syn PFFs, treatment with SAR446159 reduced the spread of aSyn pathology as measured by phosphorylated α-Syn staining and lessened the severity of motor phenotypes. Additionally, in 9-month-old transgenic mice overexpressing α-Syn (mThy1-α-Syn, Line 61), repeated treatment with SAR446159 reduced markers of α-Syn aggregation in the brain. SAR446159 had significantly higher brain and CSF penetration over a sustained period than its monospecific counterpart (1E4) in rats and monkeys. The binding properties of SAR446159 combined with its brain-shuttle technology make it a potent, next-generation immunotherapeutic for treating synucleinopathies.

## Introduction

Synucleinopathies are neurodegenerative diseases characterized by the pathological misfolding and aggregation of alpha-synuclein (α-Syn). This group of diseases includes Parkinson’s disease (PD), dementia with Lewy bodies (DLB), and multiple system atrophy (MSA). In PD, genetic studies have linked missense mutations and duplications of the gene encoding for α-Syn (*SNCA*) with autosomal dominant forms of disease^1,2^. Likewise, in sporadic PD, *SNCA* remains a strong risk locus for developing disease^3^. Though the initial conditions that first trigger α-Syn aggregation are not fully understood, once the protein converts to an aggregated, fibrillar form, it is able to template the aggregation of other monomeric α-Syn molecules^4,5^. The prion-like behavior of α-Syn aggregates is thought to underlie the spreading of α-Syn pathology throughout the brain as the disease progresses.

Prion-like spread of α-Syn aggregates was demonstrated in cell culture systems as well as in animal models. Treatment of either primary mouse neurons or human cells with α- Syn PFFs induced the conversion of the endogenous α-Syn into misfolded aggregates^4,5^. Similarly, PFFs injected into α-Syn overexpressing or even naïve mice, produced a progressive pathology that spread from the site of injection to sites connected to that region^6,7^. More recently, this templating phenomenon was taken advantage of to develop biomarkers such as real-time quaking-induced conversion (RT- QuiC)^8^ and protein-misfolding by cyclic amplification (PMCA)^9^ to evaluate whether patients carry aggregated synuclein in their cerebrospinal fluid (CSF), providing a conclusive diagnosis of synucleinopathy. Recent analysis of early and prodromal PD patient CSF has demonstrated that the presence of aggregated α-Syn is one of the earliest markers of disease, in many cases preceding the onset of clinical symptoms^10^. If these spreading-competent aggregates could be intercepted and cleared, it could potentially slow or stop the progression of disease.

The use of therapeutic monoclonal antibodies to neutralize pathogenic factors in humans has transformed the treatment of many diseases, from cancer to autoimmune disorders^11^. Though the use of biologics for treating neurodegenerative disease has been much more limited, the impact of antibody-based treatments in multiple sclerosis^12–14^ and the very recent advances in antibodies targeting amyloid beta for the treatment of Alzheimer’s disease demonstrate the untapped potential for this modality in an area of great unmet need^15,16^. The challenges faced in developing biologics for central nervous system (CNS) diseases are two-fold. First, the exact nature of the toxic species of protein aggregates has been highly debated^17^. While five anti-α-Syn antibodies have been in the clinic to date, targeting different epitopes and with varying selectivity for aggregates over monomers^18–22^, only three are still under active clinical development, with limited, but potentially promising efficacy^20,21,23^. Second, delivery of large-molecule therapeutics to the brain is hampered by the BBB. For a typical antibody, only ∼0.1- 0.5% of the infused drug substance reaches the brain^24–27^.

Receptor-mediated transcytosis (RMT) represents a pivotal strategy for enhancing the delivery of therapeutic agents, including antibodies, across the BBB, a significant impediment to effective CNS drug delivery^28^. This physiological process leverages the natural cellular mechanism of transporting molecules across endothelial cells of the BBB via specific receptor-ligand interactions. By engineering therapeutic antibodies or their carriers to target these endogenous receptors, such as the transferrin receptor (TfR), insulin receptor, insulin-like growth factor 1 receptor (IGF1R), CD98 heavy chain, or low-density lipoprotein receptor-related protein-1 (LRP1), RMT facilitates the transport of these therapeutics from the blood into the brain parenchyma^26,27,29–31^. Understanding and optimizing RMT is a frontier in neuropharmacology, combining molecular biology, engineering, and clinical medicine to overcome one of the most challenging barriers to drug delivery.

Here we demonstrate the utility of an α-Syn aggregate targeting human IgG1 antibody (1E4) coupled in the Fc domain with an engineered IGF1R targeting scFv, acting as a brain shuttle (Grabody B) – SAR446159 (also known as ABL301) (Figure 1A). Currently, a clinical phase 1 trial of SAR446159 is ongoing to evaluate the safety, tolerability, and pharmacokinetics (PK) profile following intravenous (IV) single ascending dose and multiple ascending dose administration in healthy adult participants (NCT05756920). This molecule binds to the C-terminal portion of α-Syn (Figure 1B, red) and is highly selective for aggregates over monomers. It has enhanced brain exposure compared to its counterpart lacking the brain shuttle and, consequently, improved activity in preclinical models of synucleinopathies.

**Figure 1.**
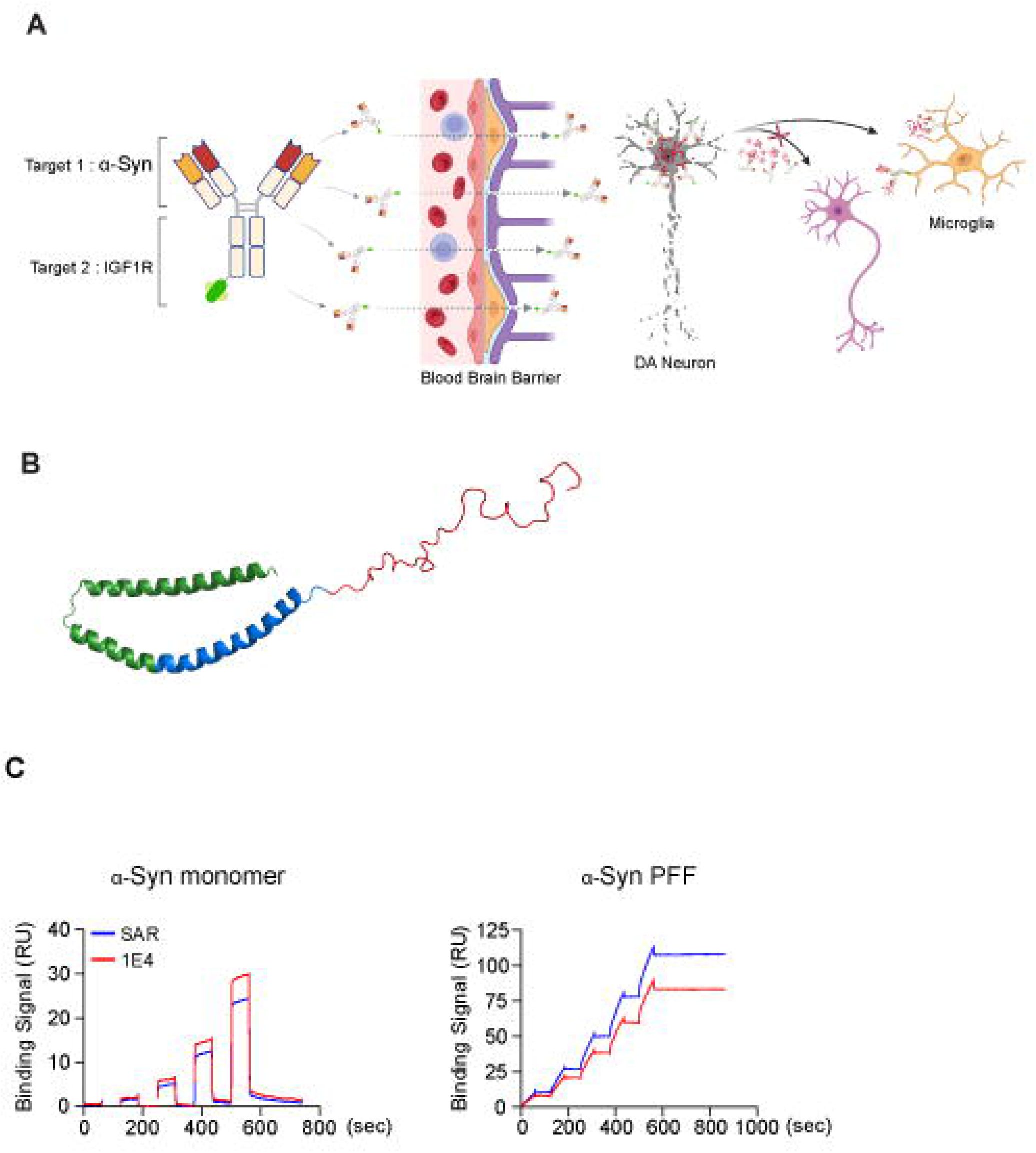
A) Schematic diagram of SAR446159 and its mechanism of action. Images were generated with the assistance of Biorender. B) The structure of aSyn, showing the the N-terminal (green) amphipathic region, the hydrophobic NAC (blue) region, and the acidic C-terminal (red) region (PDB: 1XQ8). C) Sensograms of SAR446159 and 1E4 binding to monomeric and aggregated aSyn.

## Results

### SAR446159 preferentially binds aggregated α-Syn

The affinity of the antibodies to monomeric and aggregated α-Syn were determined by surface plasmon resonance (SPR). Both 1E4 and SAR446159 bound preferentially to fibril conformations of α-Syn (Figure 1C, Table 1). Compared to the weak binding to α- Syn monomers in the hundreds of nanomolar (Figure 1C Table 1), both molecules displayed a five order of magnitude (>100,000) increased affinity for the PFF conformation of α-Syn and approached the limit of detection for SPR (sub-picomolar).

**Table 1.**
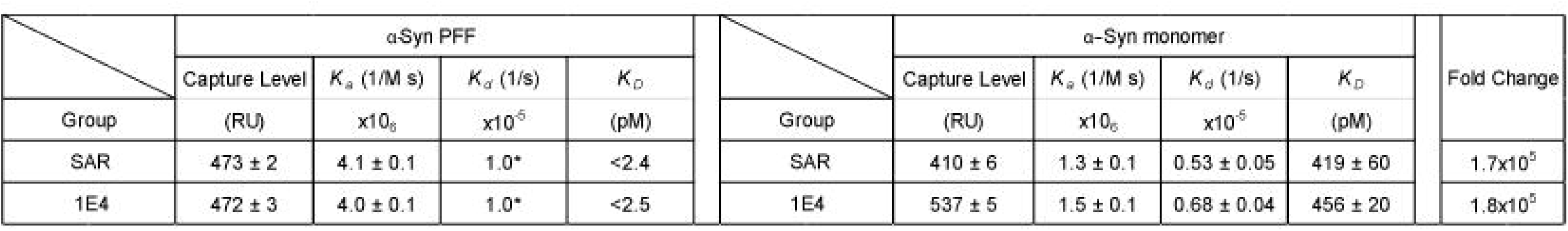
Kinetic binding parameters of SAR446159 and 1E4 with α-Syn PFF or monomers. The off rate (kd) for the binding of the α-Syn PFF was below the limit of detection of the instrument and was fixed to the lowest limit for the dissociation time used in the experiment (1.0×10^-5^ 1/s). The molarity of α-Syn aggregates was based on a particle size of 1 MDa. As a result, the binding affinities (KD) are estimates. The preferential binding of both molecules to α-Syn PFF over the monomer resulted in a five order of magnitude difference in affinity. All values are the average of n = 3.

### SAR446159 can detect pathological α-Syn in PD and MSA brain tissue

Cryogenic electron microscopy (Cryo-EM) imaging of α-Syn aggregates from patients with different synucleinopathies has revealed that the structures of the aggregates differ between diseases^32^. For a therapeutic to be effective across a spectrum of synucleinopathies, it must be able to engage the pathological form of α-Syn regardless of the structure of the aggregate. To test whether SAR446159 meets this criterion, we tested whether 1E4 could engage its target in different disease subtypes by staining human brain tissue derived from patients with PD (Figure 2, top) and MSA (Figure 2, bottom). In both sets of tissues, 1E4 was able to stain Lewy-body (LB) and Lewy-neurite (LN) structures (marked with red and blue arrows respectively) and a few dispersed intracellular α-Syn aggregates (marked with black arrows) across multiple brain regions, including the cerebral cortex, subcortical white matter, substantia nigra, substantia nigra pars compacta, and the midbrain.

**Figure 2.**
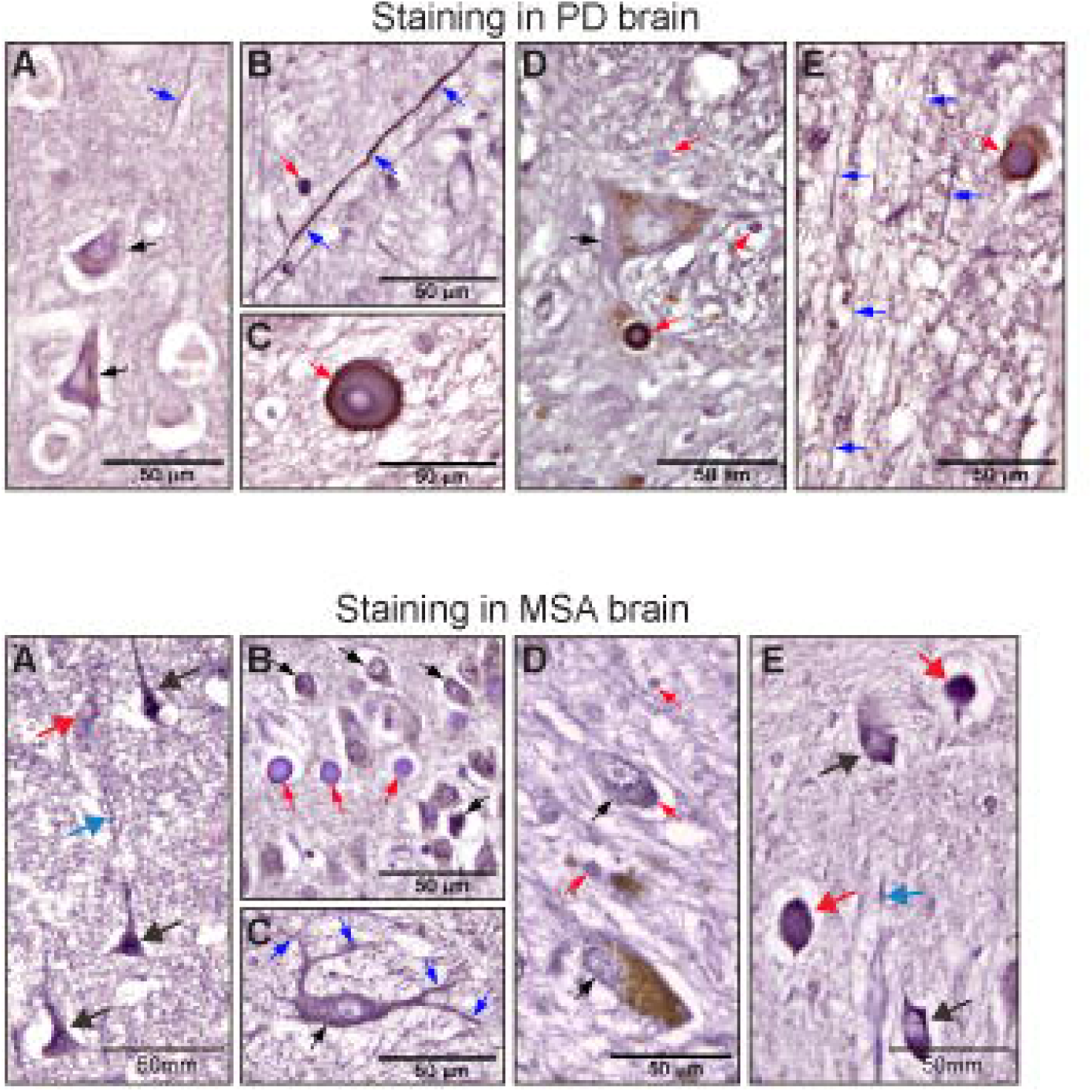
Staining of α-Syn pathology in postmortem PD and MSA patient brain tissues. In the PD patient staining, the brain regions are A) cerebral cortex, B) subcortical white matter, C) substantia nigra, D) substantia nigra pars compacta, E) midbrain. In the MSA patient staining, the brain regions are A) cerebral cortex, B) hippocampus, C) midbrain, D) substantia nigra pars compacta, E) olivary nucleus in brainstem. Black arrows are α- Syn-expressing neurons (PD) or cytoplasmic glial inclusion (MSA); red arrows, Lewy bodies; blue arrows, Lewy neurites.

### SAR446159 blocked neuronal uptake of α-Syn PFFs and clearance by microglia

We used pHrodo (a pH sensitive fluorophore) labelled α-Syn to examine how SAR446159 affected uptake of α-Syn PFFs into human induced pluripotent stem cell (iPSC) derived dopaminergic neurons and iPSC-derived microglia. In cultures without antibody co-treatment, dopaminergic neurons internalized α-Syn as visualized by an increase in the fluorescence of the pH sensitive dye in endosomes and lysosomes (Figure 3A, ‘No SAR’). This signal was significantly decreased by co-treatment with SAR446159 (Figure 3A, ‘+SAR’ at upper right). In microglia, the opposite outcome was observed. Co-treatment of microglia with SAR446159 and α-Syn PFFs increased uptake compared to microglia treated with α-Syn PFFs alone (Figure 3A) in line with the robust effector functions of the human IgG1 isotype. The SAR446159-induced microglial clearance of PFFs was mediated through its interaction with Fc gamma receptors (FcγR). Amino acid replacement of SAR446159 at its Fc (asparagine at 297 to alanine), a change known to reduce the interaction of the FcγR,^33^ abolished the uptake activity (Figure S1).

**Figure 3.**
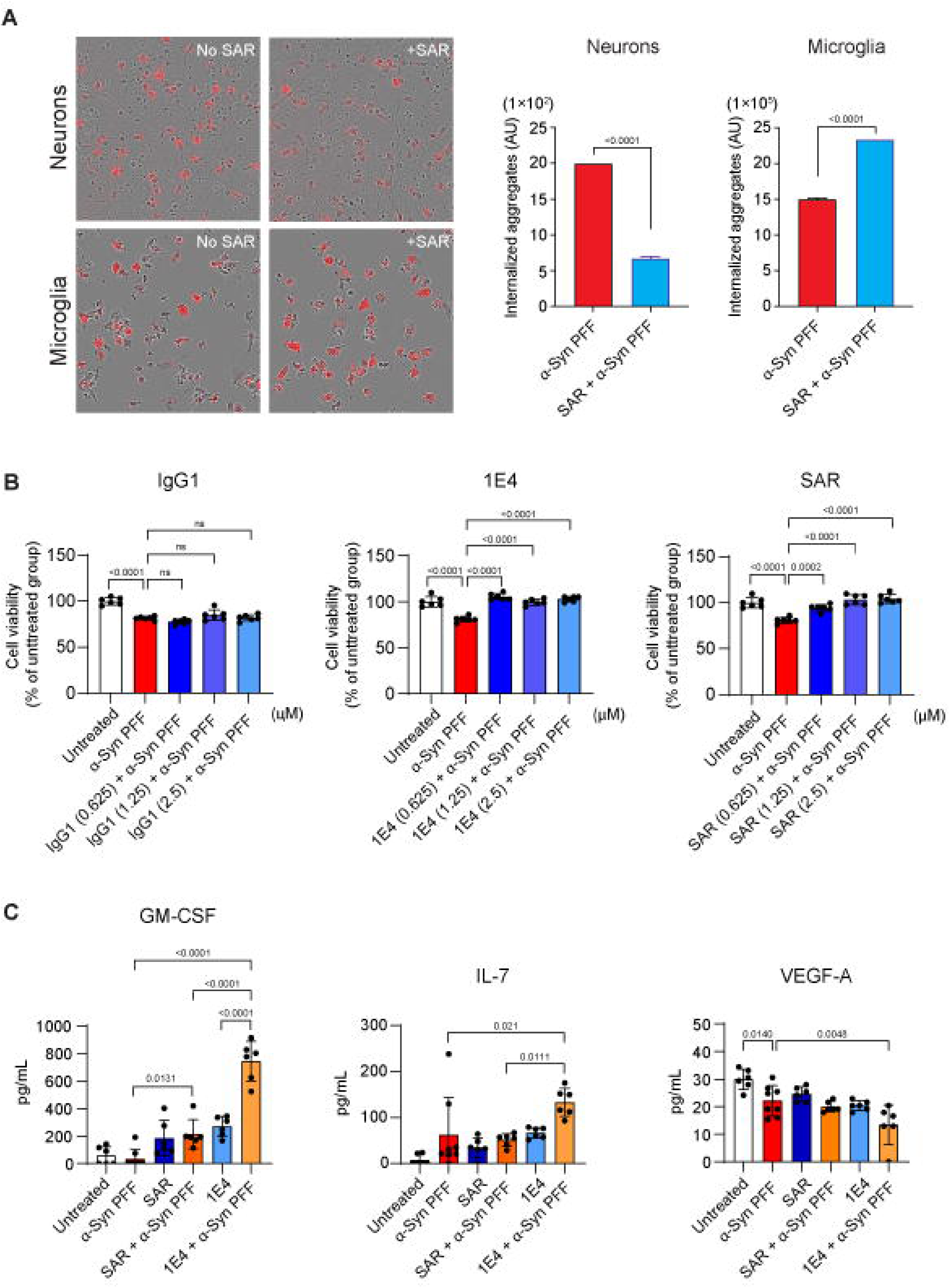
A) Human iPSC-derived dopaminergic neurons or human microglia treated with α-Syn PFFs conjugated to pHrodo (5 μg/mL), with (+SAR 10 μg/mL) or without SAR446159 (No SAR). The data were quantified at 24 hours and normalized for relative confluence. B) Mouse cortical neuron survival following high-dose PFF treatment. One-way ANOVA, followed by Dunnett’s multiple comparison’s test. C) Cytokine secretion from human iPSC-derived dopaminergic neuron-astrocyte-microglia tricultures following treatment with α-Syn PFFs with or without antibodies. PFF = 10 μg/ml, antibodies = 10 μg/ml. One-way ANOVA, Sidak correction for multiple comparisons, error bars represent S.D.

Considering that SAR446159 could block uptake of α-Syn aggregates by neurons, we tested whether it could mitigate cell death induced by large concentrations of α-Syn PFFs. Primary cortical neurons isolated from embryonic day 18 (E18) mice were cultured *in vitro* for 7 days and then treated with α-Syn PFFs and antibodies (Figure 3B). Treatment with α-Syn PFFs produced a modest loss of cells, which was not corrected by the co-addition of a non-binding, isotype control antibody (IgG1). In contrast, both 1E4 and SAR446159 were strongly protective against PFF-induced cytotoxicity at all doses tested (Figure 3B).

### Inflammatory responses to SAR446159 and α-Syn PFFs

Given the differences that we observed when cells were treated with the PFFs and antibodies, we sought to characterize how this influenced neuroinflammation. Standard monocultures cannot recapitulate the cross-cell type interactions that are essential to the function of the CNS. Previously characterized human iPSC-derived tri-culture systems, containing astrocytes, neurons, and microglia, were able to model neuroinflammation relevant to PD^34^. We stimulated tri-cultures with 1E4 or SAR446159 in combination with α-Syn PFFs and measured inflammatory factors in culture supernatants using a MILLIPLEX panel. To our surprise, there was a numerical increase in cytokines in response to α-Syn, but none of these changes were statistically significant (Figure 3C, Figure S2). Vascular endothelial growth factor A (VEGF-A) was significantly decreased in response to α-Syn PFF treatment. The most significant cytokines increased were in response to the 1E4 antibody co-treated with α-Syn PFFs, which induced interleukin-3 (IL-3), interleukin-7 (IL-7), macrophage inflammatory protein-1 alpha (MIP-1α), and granulocyte-macrophage colony-stimulating factor (GM-CSF). Notably, SAR446159 co-treatment with α-Syn PFF induced smaller changes in GM-CSF and IL-7 level than 1E4. VEGF-A and interleukin-6 (IL-6) were reduced in the same conditions that increased IL-3, IL-7, and GM-CSF (Figure S2). However, 1E4 by itself significantly reduced VEGF-A whereas SAR446159 did not. Treatment with 1E4 and α-Syn PFFs also more significantly reduced VEGF-A than SAR446159 and α-Syn PFFs. Treatment with α-Syn reduced levels of macrophage-derived chemokine (MDC). Interleukin-8 (IL-8) levels were reduced by individual treatment with α-Syn PFFs or antibodies, but antibody-PFF co-treatment increased IL-8 relative to antibodies alone (Figure 3C, Figure S2).

### SAR446159 exhibits improved brain and CSF exposure over 1E4

Grabody B, the IGF1R-binding moiety of SAR446159 is equally reactive to rat, non-human primate (NHP), and human IGF1R^29^. Therefore, transgenic animals humanized for the brain-shuttle receptor were not necessary to evaluate tissue exposure, as is the case for in vivo studies with many brain-shuttled antibodies^27,35^. Rats were infused intravenously molar equivalent doses of 1E4 or SAR446159 (30 mg/kg and 35.1 mg/kg, respectively), after which brain, CSF, and serum exposures were evaluated as a function of time. While the serum levels of both antibodies were nearly superimposable over time, SAR446159 had superior CSF and brain exposure at all time points measured except 168 hours (Figure 4A, Table 2).

**Figure 4.**
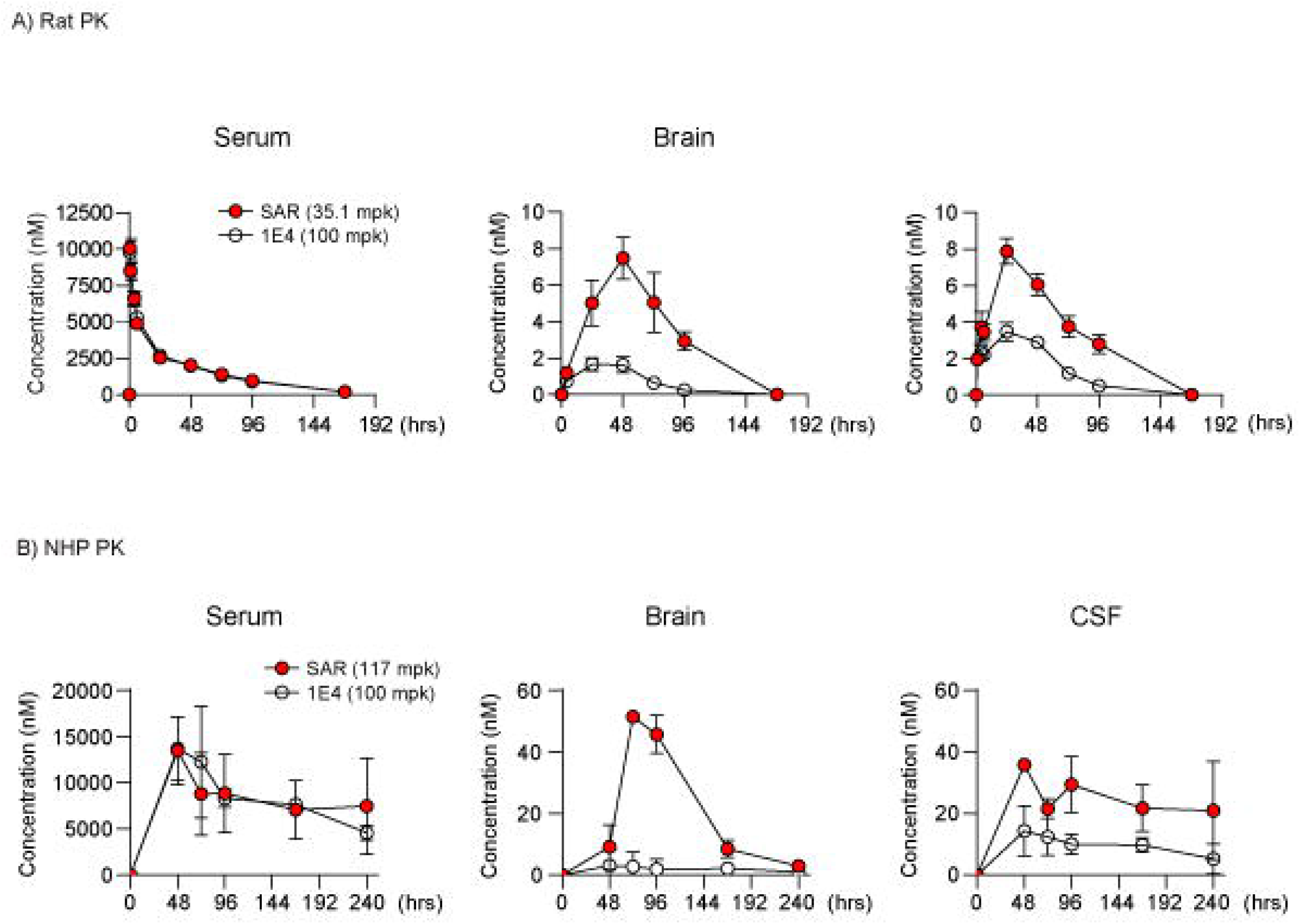
Antibody exposure of SAR446159 (red circles) and 1E4 (white circles) in the serum, brain, and CSF as a function of time following a single IV dose in A) rats (n = 5-7 animals for serum and CSF, n = 3-4 animals for brain) and B) non-human primates (NHPs, n = 3 animals). Error bars represent S.D.

**Table 2.** AUC_0-τ_ values estimated from non-compartmental analysis of mean serum, CSF and brain profiles following an intravenous bolus administration of 1E4 or SAR446159 to naïve rats. AUC_last_ represents area under the plasma concentration time curve from time zero to the last measurement.

|  | Unit \ Group | 1E4 | SAR |
| --- | --- | --- | --- |
| Serum AUC<sub>last</sub> | hr*mg/mL | 40.5 | 47.8 |
| CSF AUC<sub>last</sub> | hr*mg/mL | 0.03 | 0.08 |
| Brain AUC<sub>last</sub> | hr*mg/mL | 0.015 | 0.08 |
| CSF AUC<sub>last</sub>/Serum AUC<sub>last</sub> | % | 0.08 | 0.17 |
| Brain AUC<sub>last</sub>/Serum AUC<sub>last</sub> | % | 0.04 | 0.18 |

AUC<sub>last</sub> represents area under the plasma concentration time curve from time 0 to last measured concentration

Next, both antibodies were evaluated in NHPs. SAR446159 was administered at three doses (11.7 mg/kg, 35.1 mg/kg, 117 mg/kg), while 1E4 was measured at two equimolar doses (30 mg/kg and 100 mg/kg) (Figure 4B and Figure S2). Compounds were administered intravenously, and the serum, CSF, and brain of these animals were sampled at five time points over a period of up to 10 days (240 hours). Like the observations in rats, the serum profiles of both compounds were similar, while the brain and CSF exposure of SAR446159 was significantly greater than that of 1E4 (Table 3). Notably, the absolute differences in serum/brain and serum/CSF ratios between the two molecules were greater in NHPs than in rats.

**Table 3.** AUC_0-τ_ values estimated from non-compartmental analysis of mean serum, CSF and brain profiles following an IV bolus administration of 1E4 and SAR446159 to naïve NHPs. AUC_last_ represents area under the plasma concentration time curve from time zero to the last measurement.

|  | Group<br>Unit | 1E4 |  |  | SAR |  |  |
| --- | --- | --- | --- | --- | --- | --- | --- |
|  |  | (mg/kg) |  |  | (mg/kg) |  |  |
|  | Dose | 30 | 100 |  | 11.7 | 35.1 | 117 |
| Serum AUClast | hr*mg/mL | 80.3 | 336 |  | 31.7 | 77.6 | 458 |
| CSF AUClast | hr*mg/mL | 0.10 | 0.32 |  | 0.14 | 0.31 | 0.95 |
| Brain AUClast | hr*mg/mL | 0.01 | 0.07 |  | 0.06 | 0.20 | 0.77 |
| CSF AUClast/Serum AUClast | % | 0.12 | 0.09 |  | 0.44 | 0.40 | 0.21 |
| Brain AUClast/Serum AUClast | % | 0.02 | 0.02 |  | 0.18 | 0.26 | 0.17 |

AUClast: area under the serum concentration time curve from time 0 to last measured concentration

### SAR446159 reduced protein aggregate burden in aged mice overexpressing human α-Syn

To test SAR446159 in a rescue setting, we treated 9-month-old mThy-1 human α-Syn transgenic mice with either a control human IgG (hIgG), 1E4 or SAR446159. Animals were treated with a total of 4 doses at 3-day intervals with a dose of 50 mg/kg for 1E4 and control, and 58.5 mg/kg for SAR446159, the molar equivalent doses for these monoclonal antibodies. Forty-eight hours after the last dosing, antibody penetration into the brain parenchyma was quantified by measuring the percent area of human IgG (hIgG) immunoreactivity (red) (Figure 5A). The 1E4 group did not differ in exposure from the hIgG group in the cerebral cortex, amygdala, and substantia nigra pars compacta (SNpc). The SAR446159 group had a statistically significant increase in percent area of human IgG with an approximately 3-fold increase in the cerebral cortex and amygdala and an approximately 5-fold increase in the SNpc compared to hIgG (Figure 5A). Compared to the 1E4 group, the SAR446159 group also showed a significant increase in percent area of human IgG signals with a 3.3-fold increase in the cerebral cortex and amygdala and a 2.7-fold increase in the SNpc (Figure 5A, Figure S4A).

**Figure 5.**
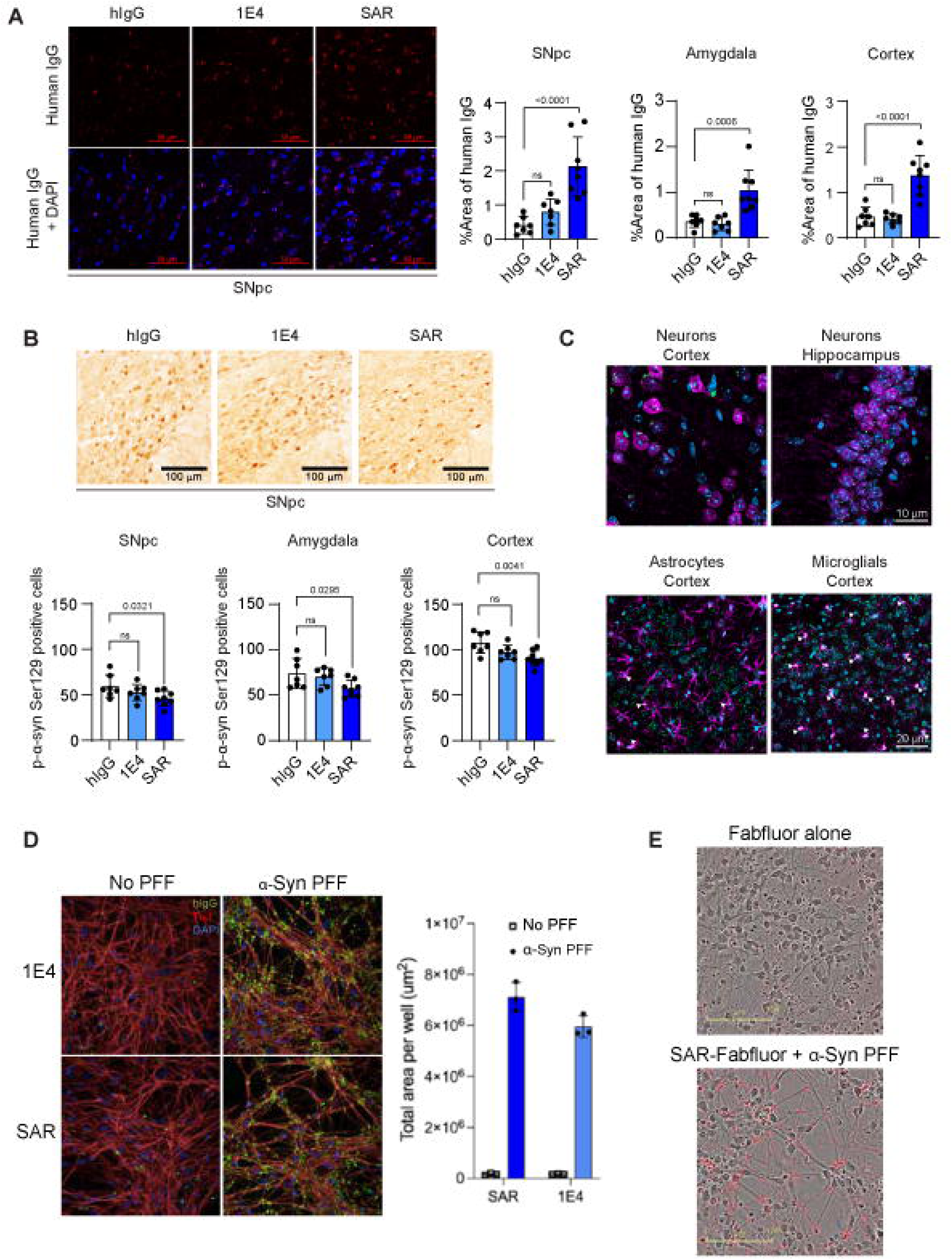
A) Human IgG staining of brain tissue from 9-month-old mThy-1 human α-Syn tg mice treated with hIgG, 1E4, or SAR446159. Representative images from the SNpc are shown. N=7 for hIgG treatment and n=8 for 1E4 and SAR446159 treatment. B) Immunostaining for pSer129 α-Syn positivity in SNpc. For each animal, average values measured from 2 fields were used. C) Co-localization of SAR446159 (hIgG stain, green) with neurons (NeuN, purple), astrocytes (GFAP, purple), and microglia (Iba1, white). DAPI was used to stain nuclei (blue). D) α-Syn-PFF enhanced localization of anti-α-Syn antibodies to neurons. DIV10 tricultures were treated with 10 μg/ml 1E4 or SAR446159 with or without 2 μg/ml α-Syn-PFF for 48 hours. Cells were fixed and IF stained against human IgG (green), neuronal marker Tuj1 (red) and DAPI (blue). E) Live cell imaging of iPSC derived dopaminergic neurons treated with Fabfluor (red when internalized) or SAR446159 labeled with Fabfluor (red when internalized, 3 μg/ml) and α-Syn PFFs (2.5 μg/ml). Statistics: all statistical analyses involved a one-way ANOVA followed by Dunnett’s multiple comparisons test. Error bars represent S.D.

Next, changes in phosphorylated α-Syn at serine 129 (p-α-Syn) expression was quantified by measuring the number of cells expressing p-α-Syn (Figure 5B). There was no difference between the 1E4 group and the hIgG group in the cerebral cortex, amygdala, and SNpc on p-α-Syn expression as measured by the number of cells with p- α-Syn staining. Despite the short duration of the treatment and the significant pre-existing synuclein pathology of the animals, SAR446159 treatment resulted in modest but statistically significant reduction in the number of p-α-Syn expressing cells in the cerebral cortex, amygdala, and SNpc compared to the control hIgG group (Figure 5B and Figure S4B).

### Cellular localization of SAR446159 is dependent on α-Syn aggregates

To understand the cellular localization of the antibody we immunostained brain tissues from the same animals in the section above to identify the cell types most often co-localized with SAR446159. Co-staining against hIgG (i.e. labeling SAR446159) and cellular markers clearly presented hIgG signals near or coinciding with Iba1, a microglial marker (hIgG: green; Iba1: white). hIgG signals also colocalized with NeuN, a neuronal marker (hIgG: green; NeuN: magenta), but not with GFAP, an astrocytic marker (hIgG: green; GFAP: magenta; Figure 5C).

Next, we examined the cellular localization of SAR446159 and 1E4 in human iPSC derived cell cultures. In tricultures without α-Syn PFF treatment, there was little or no co-localization between neuronal cell markers and fluorescently labeled SAR446159 or 1E4 (hIgG: green, Tuj1: red, Figure 5D). When cells were treated with both α-Syn PFFs and antibodies, both SAR446159 and 1E4 strongly co-localized with neuronal markers. To distinguish between localization to the cell surface and the interior, we treated iPSC derived dopaminergic neuron monocultures with α-Syn PFFs and SAR446159 labeled with Fabfluor (a pH sensitive marker) (Figure 5E). Fabfluor signal could be visualized in cell bodies and neurites, suggesting that SAR446159 was internalized into cells when α-Syn aggregates were present.

### SAR446159 reduced pathology in an inducible mouse model of synucleinopathy

Injection of α-Syn PFF into the striatum of naïve mice produces robust α-Syn pathology in anatomically connected regions of the brain. This leads to the death of dopaminergic neurons in the substantia nigra, overt degeneration of dopaminergic system in striatum, which are readily detected by reduction in tyrosine hydroxylase (TH), and deficits in motor behavior^5^. This animal model allowed the testing of SAR446159 in a preventative setting. C57BL6 mice (B6C3F1/Slc, male, 3 months old) were injected intrastriatally with 5 μg of human α-Syn PFFs. Starting one week after surgery, antibodies were systemically injected weekly, and cohorts of animals were monitored up to 3 and 6 months. Due to the long duration of this study, mouse surrogates of SAR446159 (denoted mSAR) and its non-Grabody B counterpart (denoted m1E4) were used in place of the human antibodies. There were no body weight differences between the treatment cohorts, indicating that the antibodies were well tolerated. All mice had α-Syn-PFF induced pathology, as measured by 1) LB/LN-like staining in the SNpc (Figure 6A), 2) LB/LN-like staining in the amygdala (Figure 6B), 3) TH staining in the striatum (Figure 6C), 4) p-α-Syn staining in the cortex (Figure 6D), 5) loss of TH-positive neurons in the hemisphere ipsilateral to the PFF injection compared to the hemisphere contralateral to the injection (Figure 6E), and 6) motor assessments (Figure 6F).

**Figure 6.**
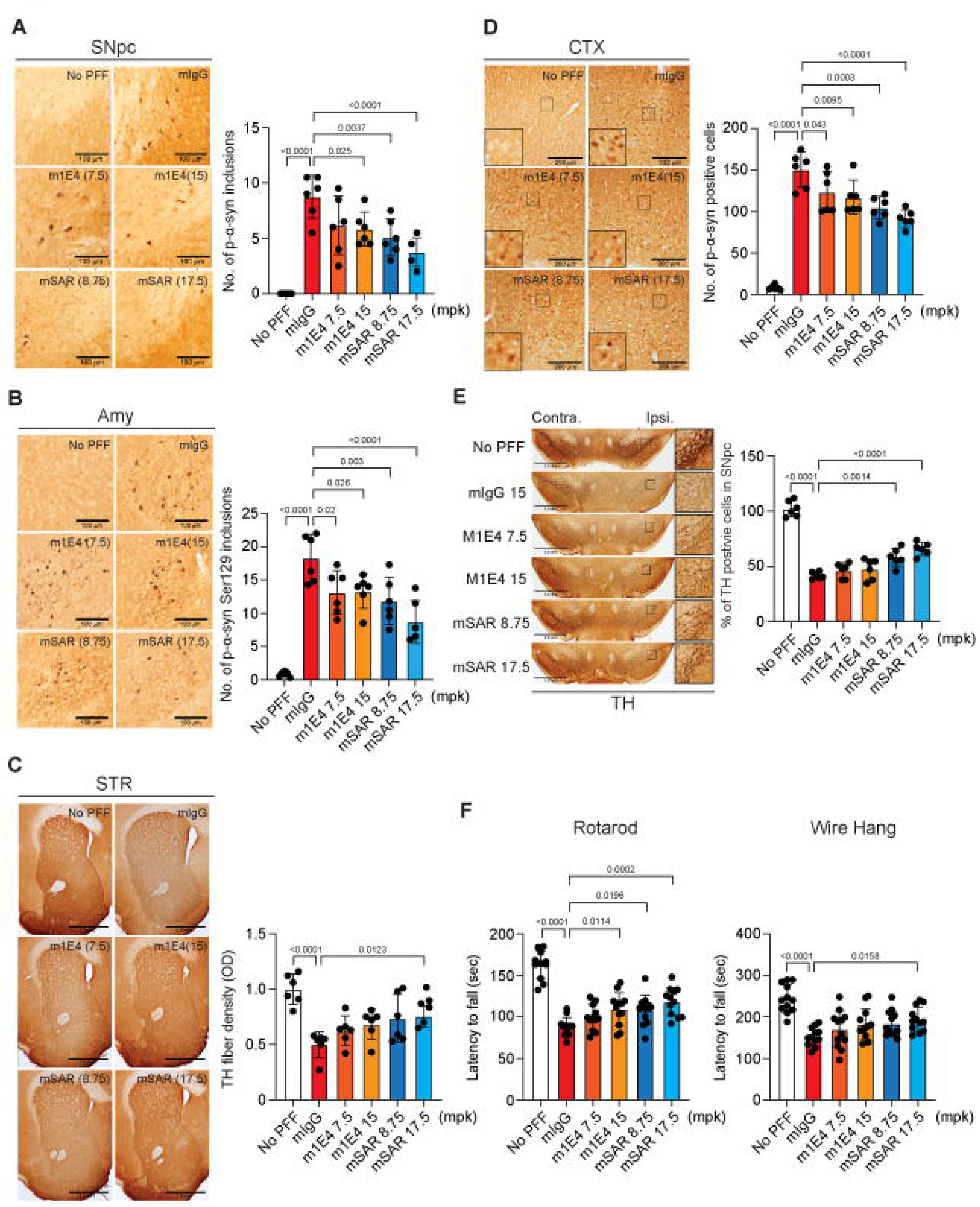
Long-term efficacy study of SAR446159. Animals were injected with PFFs at 3 months of age. One week following PFFs injection, animals were randomized and treated with weekly IV doses of antibodies. The mIgG group was treated with at dose of 15 mg/kg and the non-PFF group was treated with saline. A) LB/LN-like inclusions in the ipsilateral SNpc were counted as the average number in 4 fields (2 field per section, 2 sections per animal). B) LB/LN-like inclusions in the ipsilateral amygdala (Amy) were counted as the average number in 2 fields (1 field per section, 2 sections per animal). C) TH expression in the ipsilateral striatum (STR) was measured as optical density in 2 sections per animal. D) Cells expressing p-α-Syn in the ipsilateral cortex were counted as the average number in 6 fields (3 fields per section, 2 sections per animal). E) TH- positive cells in both the contralateral and the ipsilateral SNpc were counted as the sum of total TH-positive cells in 9 consecutive sections approximately 120 μm apart per animal. F) Latency to fall in rotarod and wire hang tests were averaged over two consecutive testing days. All experiments were performed by investigators blinded to the treatment status. Statistics: all statistical analyses involved a one-way ANOVA followed by Dunnett’s multiple comparisons test. Error bars represent S.D.

In all end points measured at either the 3-month (Figure S5) or the 6-month time point, mSAR had statistically significant improvements in more end points than m1E4. Only the animals given mSAR were protected from PFF-induced loss of TH-positive innervations in the striatum and TH^-^positive cells in the substantia nigra. Subsequently, mSAR treated animals were the most consistently protected from behavioral deficits (Figure 6F). Overall, mSAR demonstrated efficacy at both the 8.75 mg/kg and 17.5 mg/kg doses.

## Discussion

Current treatments for PD primarily address dopamine-related motor dysfunction but do not target the underlying disease pathology. The genetic link between α-Syn and PD, along with the presence of α-Syn-rich aggregates in the brains of most postmortem patients, supports the hypothesis that α-Syn plays a critical role in PD. Passive immunotherapy using α-Syn antibodies has shown efficacy in modifying synuclein pathology in animal models^18,20,36–38^. SAR446159, with its exceptionally high affinity for aggregated α-Syn, high selectivity for aggregates over monomers, and brain shuttling capabilities, has the essential characteristics of a best-in-class molecule for the treatment of synucleinopathies.

In the brain, α-Syn aggregates behave as prions, which spread along affected neuronal networks and convert monomeric α-Syn to pathological α-Syn fibrils and oligomers^6,39,40^. We hypothesized that the most effective therapeutic would specifically target these pathological particles. Because aggregated species only represent a small fraction of all synuclein particles, an aggregate-selective molecule like SAR446159 will avoid sink effects associated with the excess of non-pathogenic α-Syn monomers. Antibodies targeting the C-terminus have been shown to reduce α-Syn internalization and p-α-Syn induction *in vitro*, and lower pathology in PFF injection and mThy1-α-Syn mouse models^37,41^. Additionally, the core structure of the α-Syn folds in MSA differ from the folds found in PD and DLB patients^32^. The ability of SAR446159 to bind filaments in both MSA and PD may relate to its binding in the C-terminus outside of the core fold. Similar to other C-terminus targeted antibodies, SAR446159 binding to α-Syn PFFs blocked their internalization into iPSC-derived dopaminergic neurons. SAR446159 additionally enhanced microglial phagocytosis of α-Syn PFFs. Thus, the molecule fulfilled its dual goals of preventing α-Syn aggregate uptake by neurons and redirecting them to microglia where they can be degraded.

In addition to selectivity challenges associated with protein aggregate targeted antibodies, another potential reason for the limited efficacy of this modality is the limited brain exposure of large biologics. It is estimated that only 0.1-0.5% of a typical antibody infused by IV reaches the CNS^21,42^.To address this challenge, brain shuttles have been developed targeting antibodies for transcytosis via binding to TfR, CD98, LRP1, and IGF1R^26–31^. Such brain shuttles are already demonstrating enhanced efficacy in the clinic. For instance, trontinemab is a brain-shuttling anti-amyloid/TfR bi-specific antibody that had improved amyloid lowering activity at significantly lower doses than similar non-shuttling antibodies^43^. SAR446159 is the first brain-shuttled α-Syn targeting therapeutic to reach the clinic. It was able to partially bypass the BBB through binding IGF1R, which is more specifically expressed in the brain vasculature and parenchyma than TfR^29^. The addition of this brain shuttle increased brain exposure in both rodents and NHPs. This increased exposure was correlated with increased efficacy *in vivo.* In a human α-Syn overexpressing mouse model, four doses (administered once every three days) of SAR446159 modestly reduced aggregated α-Syn, as determined by p-α-Syn staining, whereas the non-brain-shuttled counterpart was unable to do so at molar equivalent doses. This experiment was performed in 9-month-old mice, so this observed reduction was more likely due to removal of existing aggregates rather than prevention of further buildup. Similar outcomes were achieved in a longer term, PFF-induced mouse model of synucleinopathy. While both the shuttled and non-shuttled antibodies had some benefits, at molar equivalent doses SAR446159 delivered more significant protection against neuropathology across the entire panel of outcomes.

Therapeutic monoclonal antibodies targeting pathological proteins are increasingly reaching clinical stages for treating neurodegenerative diseases. FDA approved anti-amyloid antibodies for Alzheimer’s disease show some promise in slowing disease progression but come with some significant drawbacks such as amyloid related imaging abnormalities (ARIA) and limited therapeutic efficacy^15,16^. ARIA is thought to represent edema or microhemorrhages due to removal of amyloid from the vasculature and destabilization of endothelial cells and/or inflammation^44–47^. This is not expected to be the case for α-Syn targeting antibodies, as α-Syn does not accumulate at the vasculature. Indeed, in our search of the literature, we were unable to find any case reports associating edema or microhemorrhages with a synuclein targeted antibody therapy. Despite the lack of hemorrhage risk associated with anti-synuclein antibodies, bispecific antibodies have had safety concerns derived from antibody-dependent cell-mediated cytotoxicity (ADCC) and complement-dependent cytotoxicity (CDC) in reticulocytes due to TfR targeting^48,49^. This safety liability has been linked to antibody effector function, and though silencing mutations have been introduced into therapeutic antibodies to reduce effector function, there are still liabilities relating to complement pathway activation^48–51^. Despite retaining effector function SAR446159 may avoid the undesirable reticulocyte toxicity that has been observed with TfR-shuttling antibodies by targeting IGF1R instead.

Multiple studies suggest antibody mediated amyloid clearance is at least in part microglial Fc-receptor dependent^52,53^. However, antibodies without effector function can clear tau protein aggregates in mouse models^54^, indicating that effector function is not essential for protein clearance. For SAR446159, mutations that abolished FcγR binding also lost the enhanced microglial clearance of α-Syn PFFs, indicating that this function is indeed the predominant driver of the microglial response. In addition to the ADCC and CDC risks stated above, antibody-aggregate complexes may elicit immune responses from brain-resident immune cells, especially with effector-intact antibodies. Our data indicate that such immune responses to SAR446159 are modest. In iPSC derived neuron-astrocyte-microglia tri-cultures treated with SAR446159 and α-Syn PFFs, only a handful of cytokines were altered among the broad panel examined. We showed that 1E4 antibody engagement of α-Syn PFFs triggered the release of GM-CSF, IL-3, MIP- 1a, and IL-7 by tri-cultures. The net outcome of these changes on neurodegeneration is hard to predict. For instance, MIP-1α has been implicated in autophagy disruption and increased pathological protein accumulation^55^, but extended half-life GM-CSF is being tested as a potential therapeutic in PD models^56^. Despite the reduced inflammatory functions of SAR446159 relative to 1E4, SAR446159 still induced increased iPSC-derived microglial uptake of aSyn PFFs, which we expect will be beneficial for its therapeutic application.

The well-established links between α-Syn aggregation and neurodegenerative disease support the development of targeted therapeutics. SAR446159 has a highly differentiated profile with its IGF1R targeted brain shuttle and high selectivity for α-Syn aggregates. The advantage of the brain shuttled antibody over its monospecific counterpart was the increased brain exposure, as most of the other tested properties of the molecules remained similar. Nevertheless, the increased brain exposure will enable lower effective doses in patients. The preclinical data reported here encouraged further development of SAR446159 for the treatment of synucleinopathies. A phase 1 clinical trial to evaluate the safety, tolerability, PK and PD of IV administered SAR446159 in health volunteers was initiated (NTC05656920).

## Acknowledgements

The authors would like to thank Anoushka Lotun for her assistance with cell cultures and Agnes Cheong for her assistance with the cytokine array. The authors would also like to thank James Dodge and Bradford Elmer for their careful reading and suggestions on the manuscript. The authors would like to thank Hyeran Lee, Daehae Song for their support in antibody production, Byungje Sung, Yonggyu Son for their support in quality control efforts and phagocytosis assay development, Eunsil Sung, Kyungjin Park, Jaehyun Eom for their support to generate Grabody B, Nak-Won Sohn, Jung-Won Shin for their support to interpret the human brain staining. The authors would like to thank Dao Ly, Wen Ding, Luc Tessier, Ken Chan and Sam Williamson for their support in the LC-MS-based analysis of rat and NHP PK studies.

## Author contribution

S.A., S.L., and C.K. conceived, planned, and supervised the work and wrote the manuscript. J.M. wrote the manuscript and performed *in vitro* experiments. Y.L. and O.Y. performed *in vitro* experiments. B.C.M. and Y.T. performed SPR analyses. J.M.B. and S.D. supervised the *in vitro* work and provided key reagents. L.P, R.K., S.P.S. supervised the work. D.K. cloned the antibodies used in this study, produced the antibodies, and designed the phagocytosis assay. J.A. and W.Y. designed and planned the phagocytosis assay and in vivo studies. H.Y., S.H., D.S. and H.L. produced the antibodies. S.K. and J.Y. planned the in vivo studies, and immunostained the brain sections. B.S. confirmed the quality of the antibodies prior to the in vivo study. J.J. generated PFFs used in the phagocytosis assay and in vivo studies. B.S. and R.R. provided the postmortem tissues and immunostained the human brains. J.K.S., A.H., A.T.S, C.E.D., and D.B.S. designed, planned, conducted the rat brain PK study and LC-MS-based analysis, and analyzed the NHP PK samples using the LC-MS-based method.

## Methods

### Antibodies used

1E4 was obtained from hybridoma after mouse immunization with α-Syn aggregates. Its constant domain was converted to human IgG1 to generate a chimeric version of 1E4. Subsequently, a humanized variant was finally selected after the affinity maturation.

SAR446159 was generated as an asymmetric bispecific antibody, composed of the humanized 1E4 and Grabody B with human IgG1 backbone. Grabody B, as an scFv format, was connected to the C-terminus end of a human IgG1 heavy chain using an amino acid linker. A knob-into-hole mutagenesis was introduced at its Fc to generate a monovalent bispecific antibody.

m1E4 and mSAR were generated by substitution of hIgG1 backbone of humanized 1E4 and SAR446159 as mouse IgG2a, respectively.

### α-Syn Binding Kinetics

Binding kinetics of SAR446159 and 1E4 were performed using a Biacore T200 using a CM5 Series S sensor chip (Cytiva) functionalized with streptavidin to greater than 2,000 RU (Millipore Sigma) to directly capture couple CaptureSelect Biotin Anti-LC-kappa capture antibody (Thermo Scientific) to ∼1,800 RU. The running buffer was HEPES buffered saline supplemented with 3 mM EDTA and 0.05% tween-20 (HBS-EP+). The antibodies were captured at 3 μg/mL for 60 sec at 10 μL/min to a final density of 400- 550 RU. A series of increasing antigen concentrations was injected at 30 μL/min in single cycle kinetics format starting from 500 nM (monomer α-Syn, StressMarq cat# SPR-321) or 5 nM (preformed fibrils (PFF), StressMarq, cat# SPR-322) in a 1:3 or 1:2 dilution series for 5 concentrations. The molarity of aSyn aggregates was based on a particle size of 1 MDa. The PFF was stored at −80°C until use. Prior to dilution, the PFF was rapidly thawed to room temperature in a water bath and sonicated for 10 min in 30 sec on, 30 sec off intervals. The association time was 60 sec for both antigen conformations, while the dissociation time was 180 sec and 300 sec for the monomer and PFF, respectively. The chip surface was regenerated with 10 mM glycine, pH 1.5 and 10 mM sodium hydroxide, each for 30 sec at 30 μL/min. Each binding interaction was modeled to a 1:1 fit in the Biacore Evaluation software (Cytiva) to obtain the binding kinetic parameters. All interactions were performed in triplicate and fit independently to obtain average binding parameters. The off rate of the PFF was outside the limit of detection of the instrument and was defaulted to 1×10^-5^ 1/s to obtain estimates for the binding affinities.

### Cell Culture for α-Syn and antibody uptake assays

For both human dopaminergic neuron (Fuji cat# R1088) and human microglial (Fuji cat# R1131) mono-cultures, Fujifilm protocols were followed. Briefly, dopaminergic neurons were plated at 75k cells per well on PhenoPlate 96-well plates (Revvity cat# 6055300) coated with poly-L-ornithine (Millipore cat# P4957) and laminin (Sigma cat# L2020) and maintained in Fuji dopaminergic media as instructed. Microglia were plated at 20k cells per well on PDL coated PhenoPlate 96-well plates and maintained in Fuji microglia media as instructed.

### Measuring internalization of α-Syn by pHrodo labelling

α-Syn preformed fibrils (type 1; StressMarq cat# SPR-322) were thawed at room temperature, then sonicated (Bioruptor Pico, Diagenode) at 10 °C for 10-cycles of 30 seconds on and 30 seconds off. The PFFs were then labelled according to the kit instructions (ThermoFisher cat# P36014) using 3.3 μL of reactive dye. Antibodies and labeled PFFs were added to media at 2x concentration then immediately added to cells with 50% media change for a final concentration of 10 ug/mL antibody and 5 μg/mL α- Syn-pHrodo. Plates were then live imaged in the IncuCyte SX5 and pHrodo^+^ signal was analyzed using the IncuCyte software.

### Inflammatory *in vitro* assays using human iPSC tri-cultures

Tri-culture protocols were adapted from previous Fujifilm tri-culture protocols^34^. Briefly, astrocytes were plated at 15k cells per well in PhenoPlate 96-well plates coated with PLO (Millipore-Sigma cat# P4957) and Matrigel (Corning cat# 354230), in DMEM/F12; 2% FBS (ThermoFisher cat# A3840001), and N-2 supplement (Gibco cat# 17502048). After 24-hours the full media volume was removed, and dopaminergic neurons were plated at 35k cells per well in dopaminergic neuron media. Three days after neuronal plating, microglia were plated at 10k cells per well in dopaminergic neuron media containing the microglial maintenance factors CSF1 (25 ng/mL; Peprotech cat# 300-25), IL-34 (100 ng/mL; Peprotech cat# 200-34), and TGFβ (50 ng/mL; Peprotech cat# 100- 21) through a half media change. Tri-cultures were maintained in dopaminergic neuron media with microglial maintenance factors, then treatments were added in the same maintenance media with the indicated conditions nine days after the addition of microglia. For tri-culture experiments, PFFs were added at 10 μg/ml while antibodies were at 10 μg/ml.

Partial removal of media from tri-cultures was performed (to avoid disturbing the cells) and cytokine measures were performed using a 1:5 dilution of media. All other steps were performed according to kit instructions (Millipore Sigma, cat# HCYTA-60K-PX48). For graphing and statistical purposes, with cytokines that were measured within range in most wells, cytokines under the lower limit of quantification and the lower limid of detection were set to zero. If most or all measures were out of range, results were not analyzed. M-CSF is microglial factor and was added to the tri-culture media, this represented a confounding factor, so M-CSF was not graphed. Cytokine measures and analysis were performed using Bio-Plex 200 instrument.

### Neuronal survival assay

Primary cortical neurons were prepared from timed pregnant wild-type C57BL/6JRccHsd mice at E18. Cells were counted in a hemacytometer and seeded in nutrition medium on poly-D-lysine pre-coated 24-well plates at a density of 1.8×10^5^ cells/well. Cells were cultured at 37°C; 95% humidity and 5% CO2 and a half medium exchange was performed every 3-6 days.

The treatment mixtures were prepared after 8 days-*in-vitro* (DIV8) and incubated for 1 hour at room temperature (RT) prior to addition to the cells. For that purpose, lesion agent and antibodies were combined and diluted in complete medium to obtain a 2x concentrated solution for each reagent. For treatment on DIV8, the cultivation medium was reduced to 50 μl and 50 μl of the before prepared 2x mixtures were added (70 μg/ml Alpha Synuclein A53T Mutant Preformed Fibrils (Type 1) (Stressmarq, Cat# SPR- 326) and 2x TI concentration, prepared in Neurobasal) per well. Therefore, a final concentration of 35 μg/mL (corresponds to ∼2.5 μM fibrillar or monomeric aSyn) was present on the cells, per condition n=6 wells were treated. Neurobasal alone served as untreated control.

After 48 hours of lesion, on DIV10, viability of the cultures was determined by the MTT assay using a plate-reader (570 nm). MTT solution was added to each well in a final concentration of 0.5 mg/ml. After 2 hours, the MTT containing medium was aspirated. Cells were lysed in 3% SDS and the formazan crystals were dissolved in isopropanol/HCl. Optical density was measured with a plate-reader at wavelength 570 nm. Cell survival rate was expressed as optical density (OD). Values were calculated as percent of the vehicle control.

### Antibody internalization assay with live cell monocultures

Antibodies were incubated at a ratio of 1:3 antibody to FabFluor red reagent (Sartorius cat# 4722) for 15 minutes at 37 ℃. Antibodies and α-Syn PFFs (prepared as above) were prepared in Fuji iDopaneuron medium at a 2x concentration and added to dopaminergic neuron cultures by half media change for a final concentration of 3 ug/mL antibody and 2.5 ug/mL α-Syn PFF. Cultures and pH sensitive dye were live imaged by Incucyte SX5.

### Antibody internalization assay with fixed tricultures

On DIV0, iPSC-Astrocytes (FujiFilm) were plated on PDL and laminin precoated 96-well plate at density of 15K cells per well following manufacturer’s instruction. One day later, iPSC-Dopaneurons (FujiFilm iCell DopaNeurons Kit) were plated on top of astrocyte at density at 35K cells per well. On DIV6, iPSC-microglia (BrainXell BX-0900-30) were plated at density at 15K cells per well. Every 2-3 days, half of the media was replaced with fresh media (FujiFilm Neural Base Medium 1 supplemented with 1x Neural Supplement B, 1x Nervous System Supplement, 1x iCell microglia supplement A, 1x iCell microglia supplement B, 1x iCell neural supplement C).

On DIV10, the triculture was treated with 10 μg/ml 1E4 or SAR446159 together with or without 2 μg/ml α-Syn-PFF-ATTO594 (StressMarq) for 48hr. Cells were washed with PBS, fixed with 4% PFA, permeabilized and blocked in PBS w/ 0.1% Tx-100 3% BSA and IF stained with goat anti-human IgG AF488 1:500 (Invitrogen A11013), mouse anti-Tubulin β 3 (TUBB3)-AF647 1:300 (BioLegend 801210) overnight and counter stained with DAPI. Imaging was performed using Opera Phenix and analysis was performed using Harmony 5.1 software.

### BV2 phagocytosis of α-Syn PFFs labelled with FITC

The mouse microglial cell line (BV2) was used to assess the role of Fc gamma receptor of SAR446159 in the phagocytosis of α-Syn PFFs. BV2 cells were plated on U-bottom 96 well plates at the density of 1.0×10^5^ cells/well/100 μl. FITC-labelled α-Syn PFFs were mixed with SAR446159 or SAR446159 with the NA mutation (asparagine positioned at 297 to alanine) in a tube, followed by incubation at RT for 20 minutes. The mixture was applied to the cells (total volume as 200 μl/well) and incubated at 37 ℃ for 20 hours. After a brief centrifugation, the cells were washed gently with washing buffer (PBS, pH7.4) three times, and the FITC signal was evaluated as geometric mean (gMFI) using flow cytometry (FACS Caliber, BD).

### Human brain staining

The staining of human brain sections was performed by Dr. Robert A. Rissman’s lab at University of California San Diego. Briefly, the sections were incubated at 60 ℃ for 15 minutes, followed by incubations with CitriSolvä (Deacon Labs) three times for 5 minutes each. Sections were immersed with ethanol solution serially diluted (100 %- 95 %-70 %-50 %-20 %), followed by PBS wash. After incubation with sucrose of 10% concentration for 30 minutes and multiple PBS wash, sections were blocked with 10% normal goat serum (#S1000, Vector technology) in PBS for 1 hours at room temperature. The sections were incubated with 1E4 (1:1000) at 4 ℃ overnight.

Next day, the sections were rinsed with PBS three times, and incubated with a diluent solution of a biotinylated goat anti-human secondary antibody (#BA3000, Vector technology, 1:100 in PBS) for 60 minutes at room temperature. After repeated PBS wash, ABC solution (one drop of A and B in 10 ml of PBS; ABC kit from Vector technology (PK6100)) was applied to the sections for 1 hour at room temperature. After washing with PBS three times for 5 minutes each, the 3,3’ Diaminobenzidine (DAB) solution (2 drops of Buffer solution, 4 drops of DAB, and 2 drops of H_2_O_2_ solution in 5 ml of double-distilled water (ddH2O); DAB kit from Vector technology (#SK4100)) was applied to the sections for 1 to 5 minutes, followed by rinsing with ddH2O. The sections were mounted on slides, dried for overnight, and covered with coverslip.

The sections were visualized and analyzed using a light microscope (pOlympus BH51 with DP70 CCD camera, Japan) at Alsox200 magnification at Novamedilab.

### Short term PK/PD study

mThy-1 human α-Syn transgenic (tg) mice (male, 9-month-old) were randomly divided up into three groups, with 7 animals for the hIgG control group and 8 animals in each of the treatment groups. Test articles were intraperitoneally injected a total 4 doses with 3- day intervals. Monoclonal antibodies were treated at a dose of 50 mg/kg, and SAR446159 was treated at dose of 58.5 mg/kg (molar equivalent doses for monoclonal antibodies). Mice were sacrificed and tissues were harvested 48 hours after the final treatment.

Forty-eight hours after the last dosing, mice were anesthetized with isoflurane inhalation and transcardially perfused with 20 mL of 0.05 M PBS. Brains were harvested and fixed in 4% paraformaldehyde for 24 hours at room temperature (RT), then transferred into 30% sucrose for 3 days. Brain tissue was sliced into 40 μm thick sections using a cryostat microtome (Leica, CM1850, Germany). The sections were immersed and stored at −20°C in storage solution (0.02 M phosphate buffer, sterile water, ethylene glycol, glycerin) until use.

For immunofluorescence labeling of human IgG, the sections were washed with 0.1 M PBS 2 times, followed by blocking in solution of 5% BSA and 0.3% triton x-100 for 1.5 hours at RT. The sections were incubated with Alexa Fluor 488 goat-anti-human antibody (Jackson Immuno Research, 109-545-003, 1:250) with 1% BSA at RT for 2 hours. The sections were washed in PBS three times, then were mounted on slides using VECTASHIELD mounting medium with DAPI to visualize nucleus. For immunohistochemistry against p-α-Syn, brain sections were washed twice with PBS followed by a blocking process using 3% H_2_O_2_. Detection was carried out with anti-pSer129-α-syn antibody (S129) (Abcam, ab51253, 1:500) using avidin-biotin complex system (Vectastain ABC Elite Kit, Vector Laboratories). The immunocomplexes were visualized with the chromogen 3,3’- diaminobenzidine (DAB). Images of immunofluorescence-labeled human IgG were captured with a confocal laser-scanning microscopy (Carl Zeiss, LSM 800, Germany). Images of DAB-reacted p-α-Syn were saved using a BX51 microscope (Olympus, Japan) equipped with a DP70 digital camera (Olympus, Japan).

Image analysis was performed using ImageJ v1.53 (NIH). Human IgG biodistribution was determined as the % area of immunofluorescent labeled human IgG in a defined field (160 x 160 mm^2^). Expression of p-aSyn was determined as the average number of cells expressing p-α-Syn in a defined field (320 x 320 mm^2^). For each animal, average values measured from 6 fields (3 fields per section, 2 sections per animal) in the cortex, 4 fields (2 fields per section, 2 sections. per animal) in the amygdala, and 2 fields (1 field per section, 2 sections per animal) in the substantia nigra pars compacta (SNpc) were used. Statistical analyses and calculation of p-values were performed using Prism10 (GraphPad Software); One-way ANOVA for multiple comparisons with Tukey’s post-hoc test.

For a cellular localization analysis of SAR446159, immunofluorescence staining was performed with human IgG and three cell type markers (NeuN, Iba1 and GFAP). Briefly, the floating brain sections kept at – 80 ℃ was placed on the slide with a brush. Then, the brain sections on the slide were dried out for 20 minutes at room temperature. After washing with PBS 2 times, a blocking solution (0.3% triton x-100, 10% NGS in PBS) was added for 1 hour at room temperature. Then, the sections were incubated with a primary antibody, either NeuN (Millipore, MAB377, 1:100) or Iba1 (Wako, 019-19741, 1:500) or GFAP (Dako, GA524, 1:500) for 16 hours at 4 ℃. Next day, the sections were washed with PBS for 10 minutes 3 times and incubated with a secondary antibody solution for 2 hours at room temperature. The secondary antibody combination for each primary antibody is listed; For NeuN co-staining, rabbit anti-human IgG(H+L)-FITC (Southern Biotech, 6145-02, 1:200) was mixed with Alexa Fluor 647 goat-anti-mouse antibody (Invitrogen, A32728, 1:500). For Iba1 and GFAP co-staining, goat anti-human IgG(H+L)-FITC (Southern Biotech, 2087-02, 1:200) was mixed with Alexa Fluor 647 goat anti-rabbit antibody (Invitrogen, A32733,1:500). After washing with PBS for 10 minutes 3 times, the nucleus was staining with DAPI and the sections were washed with PBS for 10 minutes 3 times. The stained sections were mounted with Aqua-Poly/mount (Polysciences, 18606-20) and covered with the coverslips. Images were achieved with a confocal laser-scanning microscopy (Carl Zeiss, LSM800 or LSM900, Germany).

### Long term PFF-induced mouse study

The long-term treatment study was led by Novamedilab Co., Ltd. (A2401-1, 184 Jungbu-daero, Giheung-gu, Yongin-si, Gyeonggi-do, Korea 1709). Animals were treated and housed at KLS Bio Inc. (Shinwon-ro,1, Suwon-si, Gyeonggi-do, Korea 16670). Behavioral analyses and tissue collection were performed at the College of Electronics and Information building, Kyung Hee University Global campus (#447-1, 1702 Deogyeong-daero, Giheung-gu, Yongin-si, Gyeonggi-do, Korea 17104). Tissue imaging and data analysis were performed at Novamedilab.

A total of 240 C57BL6 mice (B6C3F1/Slc, male, 3 months old) were used in this study; 120 mice in the 3-month treatment period and 120 mice in the 6-month treatment period. Mice were initially anesthetized with ketamine hydrochloride (50 mg/kg, intraperitoneal) and maintained in anesthesia during all surgical procedures using 2% isoflurane in O_2_-N_2_O gas (Lab Animal Anesthesia System, Matrax/SurgiVet, Matrax Medical, USA). Using an electronic drill, a skull hole (∼0.5, mm in size) was made 0.2 mm anteriorly and 2.0 mm lateral to the bregma. The α-Syn PFF (5 μg/2.5, μL per animal, StressMarq Biosciences, SPR-322) was injected into the striatum unilaterally (stereotaxic coordinates: +0.2 mm from Bregma, +2.0 mm from midline, +2.6 mm beneath the dura). The injections were performed using a 10 μL syringe (Hamilton, NV) at a rate of 0.2 μL per min using a micropump (Model 310, Kd Scientific, USA). The needle in place was maintained for >5 min after completion of α-Syn PFF injection to prevent reflux and was carefully removed. After removing the needle, the punctured hole in the skull was filled with bone wax and the skin was sutured. Animals were returned to home cage after recovery from anesthesia. Stereotactic injection of α-Syn PFF was performed to a total of 216 animals, and 24 animals (non-PFF group) underwent sham operation without α-Syn PFF injection.

One week following the α-Syn PFF injection, the animals were randomized into treatment groups. Among the test articles, monoclonal antibodies (mIgG, and m1E4) were treated at doses of 7.5, and 15 mg/kg, and the bispecific antibody (mSAR) was treated at doses of 8.75, and 17.5, mg/kg (molar equivalent doses for monoclonal antibodies). The mIgG group was treated with a dose of 15 mg/kg and the non-PFF group was treated with the same amount of saline.

For behavioral tests, animals were transported at least one week prior to the start of behavioral tests after the last dose of the 0-month and 6-month treatment periods. Mice were acclimatized to the test environment 1 week before the actual experiment. All behavioral tests were performed by an investigator blinded to the treatment conditions. To assess motor impairment and balance, the rotarod test was performed using a rotarod apparatus (3ED-Associates Rotarod Apparatus). Mice were first trained on a rotating rod accelerated from 4 rpm to 40 rpm over 5 min. After two training sessions, a test session followed once a day over 0 days. Latency to fall was recorded and averaged over three testing trials. To assess grip strength and motor coordination, the wire hang test was performed. Briefly, mice were placed on top of a stainless-steel wire mesh (1 mm wire diameter, 1 cm hole) and then turned upside down to let the animals grip the wires. The wire mesh was stirred with a shaking apparatus to gradually increase the shaking frequency from 0 Hz to 100 Hz over 1 min. After one training session, a test session followed once a day over 2 days. Trials were stopped if mice remained on the wire mesh for over 5 min (recorded as 300 second). Latency to fall was recorded and averaged over two testing trials.

The day after the behavioral testing was completed, mice were anesthetized with isoflurane inhalation and transcardially perfused with 20 mL of 0.05 M PBS. Brains were harvested and fixed in 4% paraformaldehyde for 24 hours at room temperature, then transferred into 30% sucrose for 3 days. Brain tissue was sliced into 40 μm thick sections using a cryostat microtome (Leica, 2800N, Germany). The sections were immersed and stored at −20°C in storage solution (0.02 M phosphate buffer, sterile water, ethylene glycol, glycerin) until use. Brain sections were washed twice with PBS followed by a blocking process using 1% H_2_O_2_. Immunohistochemical detection was carried out with anti-pSer129-α-Syn antibody (S129) (Abcam, ab51250, 1:500) or anti-tyrosine hydroxylase antibody (Pel-Freez, 40101, 1:1000) using avidin-biotin complex (ABC) system (Vectastain ABC Elite Kit, Vector Laboratories, Burlingame, CA). The immunocomplexes were visualized with the chromogen 3,3’-diaminobenzidine (DAB) and images were saved using an Olympus BX51 microscope equipped with a DP70 Olympus digital camera.

Image analysis was performed using ImageJ v1.53 (NIH). Cells expressing p-α-Syn in the ipsilateral cortex were counted as the average number in 6 fields (640 x 640 mm^2^, 3 fields per section, 2 sections per animal) at both the 3-month and the 6-month treatment periods. LB/LN-like inclusions in the ipsilateral amygdala were counted as the average number in 2 fields (320 x 320 mm^2^, 1 field per section, 2 sections per animal) at the 3- month treatment period and as the average number in 4 fields (320 x 320 mm^2^, 2 fields per section, 2 sections per animal) at the 6-month treatment period. LB/LN-like inclusions in the ipsilateral SNpc were counted as the average number in 2 fields (320 x 320 mm^2^, 1 field per section, 2 sections per animal) at both the 3-month and the 6- month treatment periods. TH-positive cells in both the contralateral and the ipsilateral SNpc were counted as the sum of total TH-positive cells in 9 consecutive sections approximately 120 μm apart per animal. TH expression in the ipsilateral striatum was measured as optical density (OD, gray level of TH immunoreactive intensity) and individual data were obtained by subtracting the OD background. At the 3-month treatment period, 1 section per animal and at the 6-month treatment period, 2 sections per animal were measured and the mean of both sections was calculated. The investigator who performed the image analysis blinded to the animal treatment status.

### Rat and NHP PK studies

Rat PK study was conducted and analyzed by National Research Council Canada (NRC) using the animal use protocol approved by the Institutional Animal Care Committee in accordance with the guidelines from the Canadian Council on Animal Care. Thirty-six (36) male Wistar rats (125 – 150 grams) were acclimated for a period of at least 5 days and were housed with a 12-hour light and 12-hour dark cycles at 23-24 °C with humidity of 50%. 1E4 and SAR446159 were administered via a bolus injection through the tail vein.

Blood samples were collected at a total of 9 time points (0.5, 1, 4, 6, 24, 48, 72. 96, and 168 hours) via the tail vein or via cardiac puncture (only at the terminal time point) under anesthesia with 4-5% isoflurane. CSF samples were collected at a total of 8 time points (1, 4, 6, 24, 48, 72, 96, and 168 hours) via the cisterna magna under anesthesia. Brains were collected at the terminal time points after perfusion via carotid artery with deep anesthesia (time points: 4, 24, 48, 72, 96, and 168 hours). Blood, CSF, and brain samples were frozen immediately and stored at −80 °C until the analysis.

The NHP PK study was conducted at Macaque *in vivo* facility of Atuka Inc (Suzhou, China). The facility is accredited with AAALAC and follows the Institutional Animal Care and Use Committee (IACUC)-approved protocols. Seventy-six (76) normal, previously untreated, male cynomolgus monkeys (*Macaca fascicularis,* approximately 2-3 years of age and weighing approximately 2.5 - 4.5 kg) were singly housed with a 12-hour light-dark cycle (lights on at 7 a.m., 100-200 lux), temperature 20-28 °C, relative humidity of 40-70%, ventilation frequency of 11-14 times/hour and in a room containing only animals of the same sex. The animals were acclimated to the experimental setting for a period of at least 3 weeks.

Two different doses of 1E4 (30 and 100 mg/kg) or three different doses of SAR446159 (11.7, 35.1, and 117 mg/kg) were administered intravenously. Fifteen (15) animals were assigned to each dose group except 1 animal assigned for vehicle-treated control group. Blood, CSF and brain samples were collected at 48, 72, 96, 168, and 240 hours after the administration (n = 3 per time point) with sedation with Zoletil and pentobarbital except the control animal.

Antibody levels in serum, CSF, and brain samples were quantified using targeted, multiplexed mass spectrometry in selected reaction monitoring (SRM) by the NRC. Briefly, after reduction, alkylation and trypsin digestion of the samples, peptide signatures of various antibody domains (for instance, human Fc and anti-α-Syn Fab) were identified, and used for the analysis after passing the acceptance criteria. For absolute quantification, calibration curves were created in antibody-spiked matrices. At least three quality control standards (low, mid, and high) were also prepared by independently spiking the antibodies in the appropriate matrices and run after every 5 to 10 samples. All samples were analyzed by nanoLC-SRM by injection onto a C18 or C8 PepMap™ 100 trap (Thermo Fisher, Waltham, MA) and subsequent nanoLC BEH130C18 or Atlantis^™^ dC18 column (Waters, Milford, MA) on a nanoAcquity UPLC (Waters) coupled with ESI-LTQ-XL-ETD or TSQ-Quantiva mass spectrometers (Thermo Fisher). Albumin peptide was used to identify and exclude serum-contaminated CSF samples (albumin CSF/serum ratio >0.1%).

---

**Figure S1.**
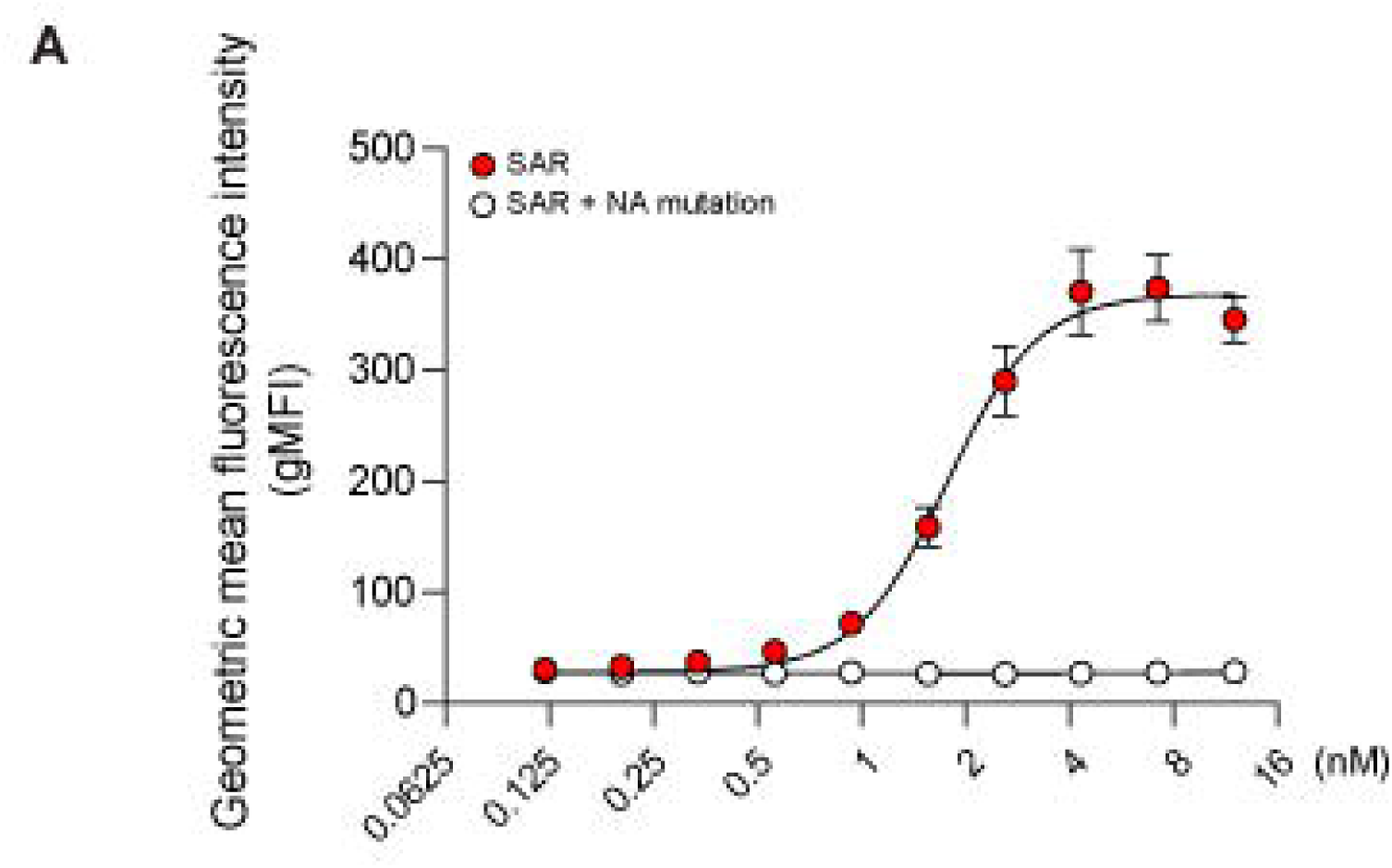
Potential role of the Fc interaction of SAR446159 with FcγR in phagocytosis of α-Syn PFFs. Note the almost no activity of SAR446159 with the NA mutation (white) in contrast to a sigmoidal increase in internalized α-Syn PFFs applied with SAR446159 (red).

**Figure S2.**
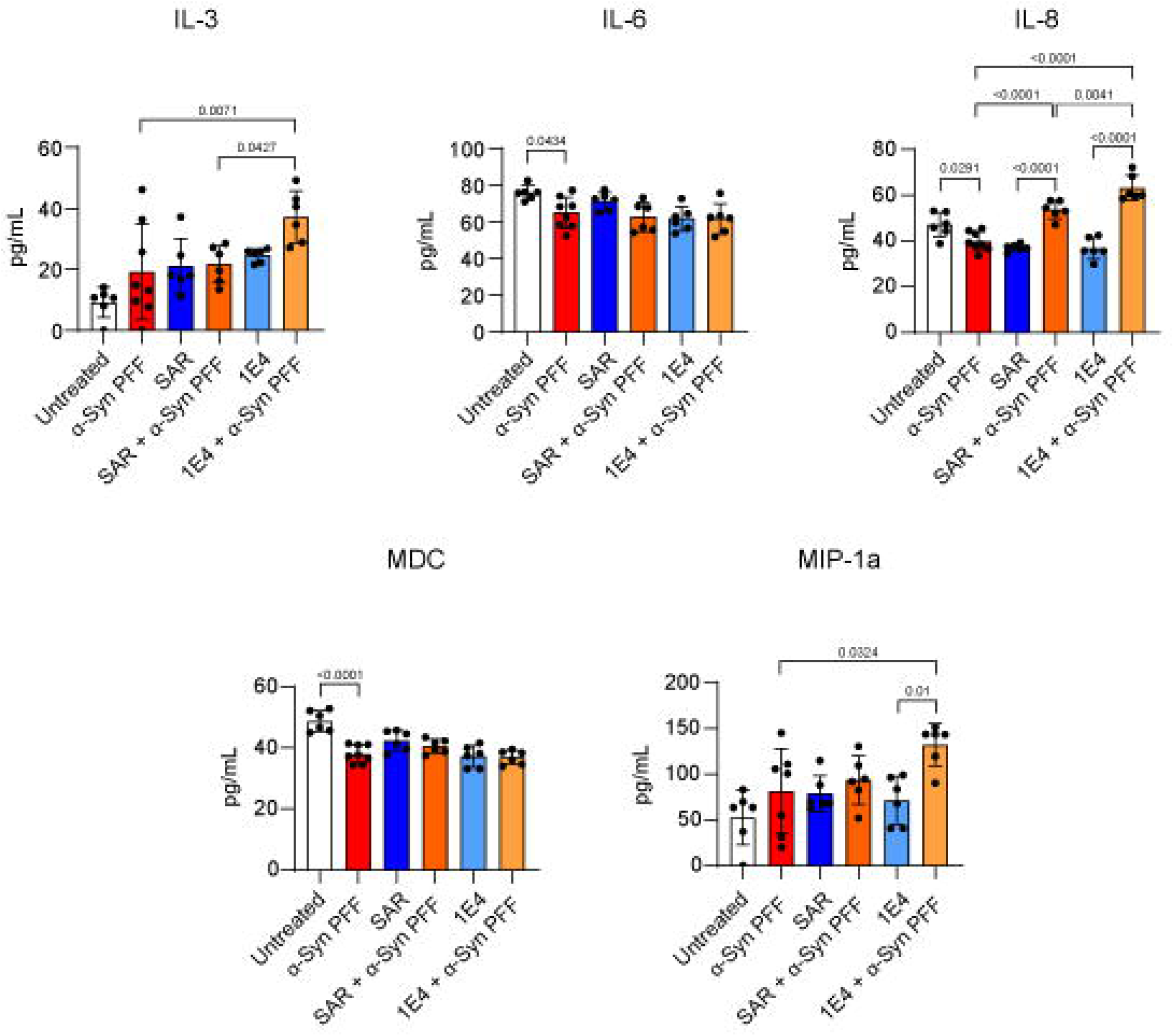
Additional cytokine secretion data from human iPSC-derived dopaminergic neuron-astrocyte-microglia tri-cultures following treatment with α-Syn PFFs with or without antibodies. PFF = 1 μg/ml, antibodies = 10 ug/ml. One-way ANOVA, Sidak correction for multiple comparisons, error bars represent S.D.

**Figure S3.**
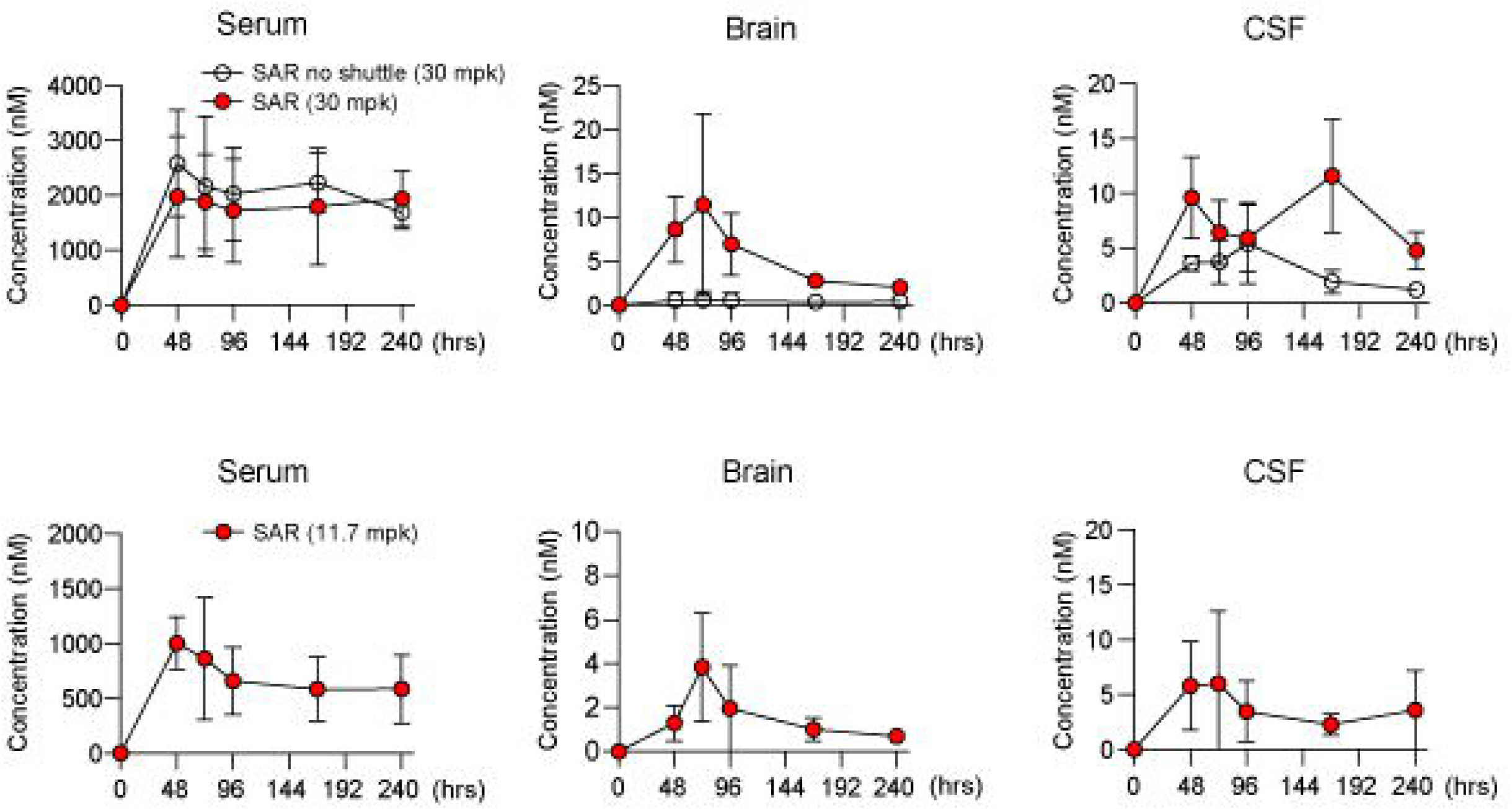
Antibody exposure of SAR446159 (red circles) and 1E4 (white circles) in the serum, brain, and CSF as a function of time following a single IV dose in NHPs (n = 3 / dose/ time point). Error bars represent S.D.

**Figure S4.**
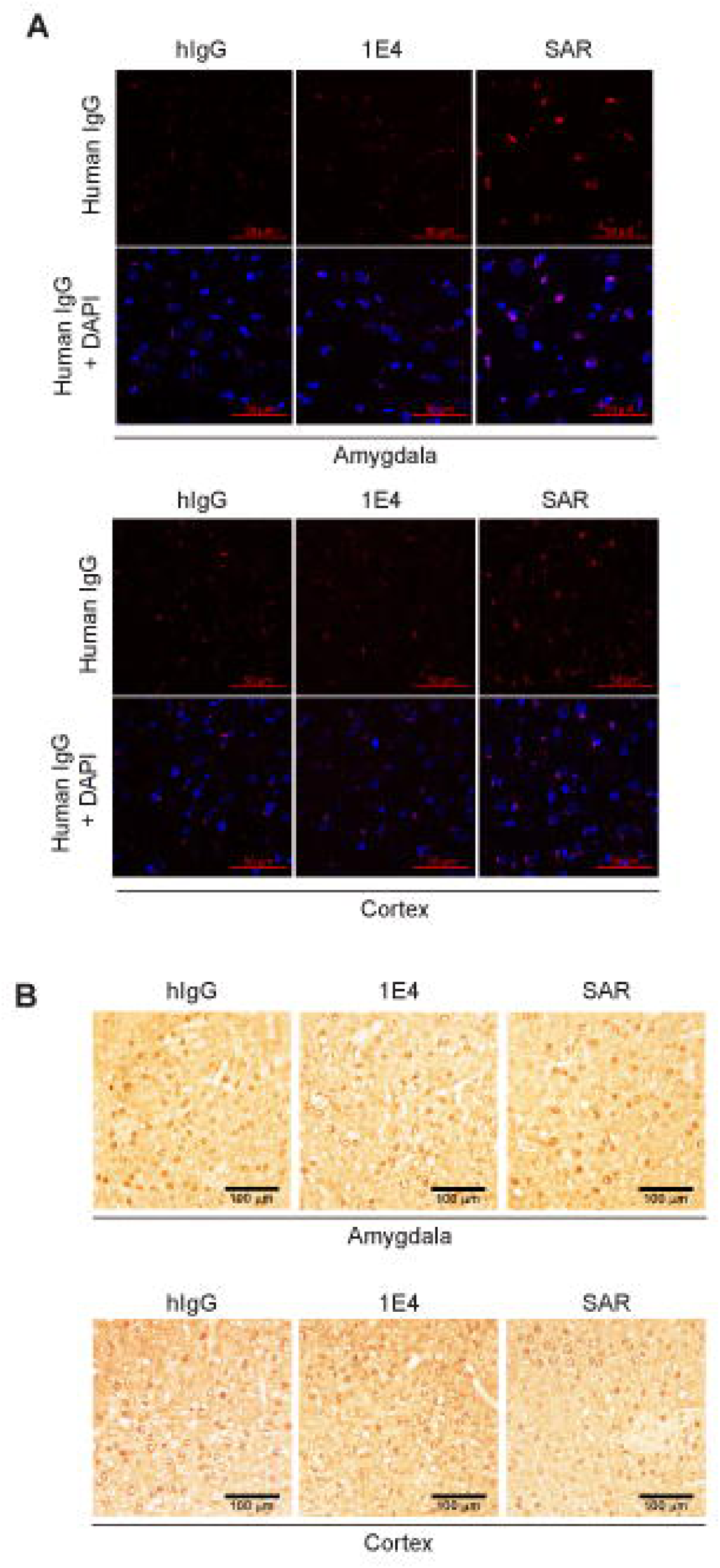
Efficacy of SAR446159 with 3-month, weekly treatment. A) LB/LN-like inclusions in the ipsilateral SNpc. B) LB/LN-like inclusions in the cortex. All experiments were performed by investigators blinded to the treatment status. Statistics: all statistical analyses involved a one-way ANOVA followed by Dunnett’s multiple comparisons test.

**Figure S5.**
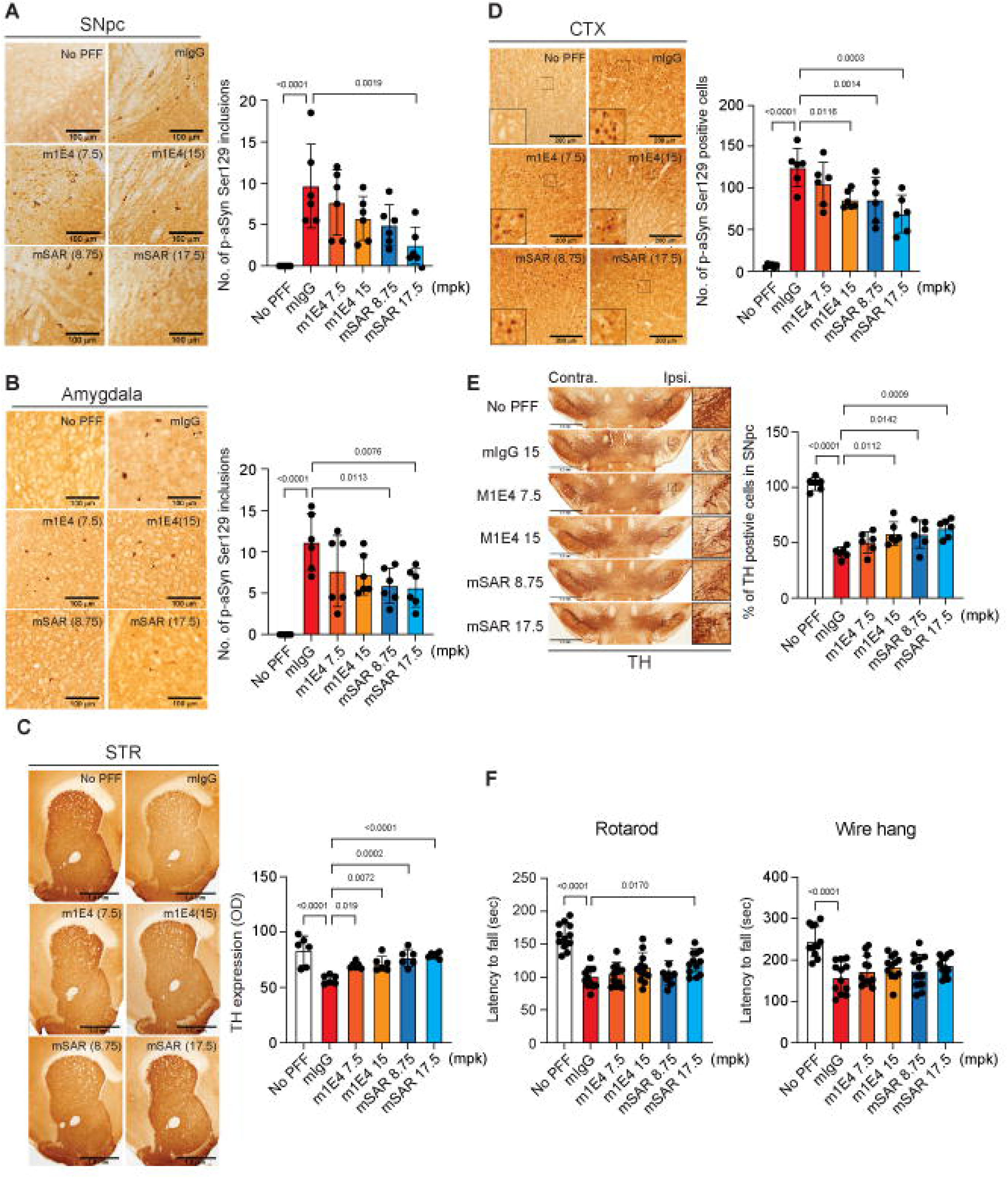
Efficacy of SAR446159 with 3-month, weekly treatment. A) LB/LN-like inclusions in the ipsilateral SNpc. B) LB/LN-like inclusions in the amygdala, C) TH expression in striatum, D) number of p-α-Syn-positive cells in cortex, E) percentage of TH positive cells in SNpc, and F) two behavioral assessments (rotarod and wire hang). All experiments were performed by investigators blinded to the treatment status. Statistics: all statistical analyses involved a one-way ANOVA followed by Dunnett’s multiple comparisons test.

